# MRI2MRI: A deep convolutional network that accurately transforms between brain MRI contrasts

**DOI:** 10.1101/289926

**Authors:** Sa Xiao, Yue Wu, Aaron Y. Lee, Ariel Rokem

## Abstract

Different brain MRI contrasts represent different tissue properties and are sensitive to different artifacts. The relationship between different contrasts is therefore complex and nonlinear. We developed a deep convolutional network that learns the mapping between different MRI contrasts. Using a publicly available dataset, we demonstrate that this algorithm accurately transforms between T1- and T2-weighted images, proton density images, time-of-flight angiograms, and diffusion MRI images. We demonstrate that these transformed images can be used to improve spatial registration between MR images of different contrasts.

## INTRODUCTION

MRI creates images that are sensitive to different aspects of the tissue, and susceptible to different imaging artifacts. The relationships between different imaging contrasts are nonlinear and spatially- and tissue-dependent (1). This poses several difficulties in the interpretation of multi-modal MRI. For example, analysis that requires accurate registration of images into the same coordinate frame currently requires the use of algorithms that can match images with different contrasts (2). While this often works well, these algorithms can be over-sensitive to large, prominent features, such as edges of the tissue, and are more error-prone when images have low SNR, or represent very different features. Moreover, a better understanding of complementary information provided in different contrasts will allow better characterization of tissue properties.

We used deep learning (DL) algorithm to learn the complex mappings between different MRI contrasts. DL has been successful in many different domains (3), and there are several previous applications of DL in biomedical imaging (4), including classification of disease states (5) and segmentation of tissue-type (6) lesions (7) and tumors (8). DL algorithms that take an image as input and generate another image as output have been used for style-transfer: retaining the content of an image, while altering its style to match the style of another image (9). For example, an image of a summer scene used to synthesize the same scene in winter (10). Here, we rely on some of the principles underlying style transfer, to transform brain MR images with one contrast into images of the same brain in other contrasts.

## METHODS

We used the IXI dataset (http://brain-development.org/ixi-dataset/): T1w, T2w, PD, MRA, and DWI (16 directions, at b-value of 1000, one b=0) for 567 are available. A convolutional neural network was trained to learn the mapping between different MRI images in a training set (n=338). We used a U-net architecture (7), with loss evaluated on “perceptual loss” (9): the activation of the first layer of a pretrained VGG16 network (11), a cost function that induces image similarity and prevents over-smoothing. Training used the Adam optimizer (learning rate: 0.0002). Implementation using Pytorch (https://pytorch.org).

A group of participants (n=79), set aside as a test set (not shown to the algorithm during training) was used to evaluate registration. We used a state-of-the-art registration algorithm (12), implemented in DIPY (13) to register DWI b=0 to corresponding T1w images, using a mutual information metric to find the best affine transformation for registration. To simulate participant motion, known rotation and translation were applied to the b=0 image and the T1w was registered to the b=0. For comparison, we also synthesized a T1w-analog from the b=0 image using the DL network and used the synthesized image with the same registration algorithm. Registration errors were calculated relative to known ground truth as mean absolute error (MAE), for translation and rotation components of the registration.

## RESULTS

The DL algorithm learns the mapping between different contrasts (Figure 1). It can be trained for either one-to-one mappings (e.g, T1w = > PD, Figure 1A, or T1w = > T2w, Figure 1B), or many-to-one mappings (e.g., T1w + T2w + PD = > MRA, Figure 1B). High accuracy is achieved by learning both mappings on the individual voxel level, as well as overall global structure. For example, the algorithm learns to disregard the ventricles in the mapping to MRA, despite the fact that the ventricles have similar pixel values in T1w, T2w and PD images to blood vessels.

**Figure 1:**
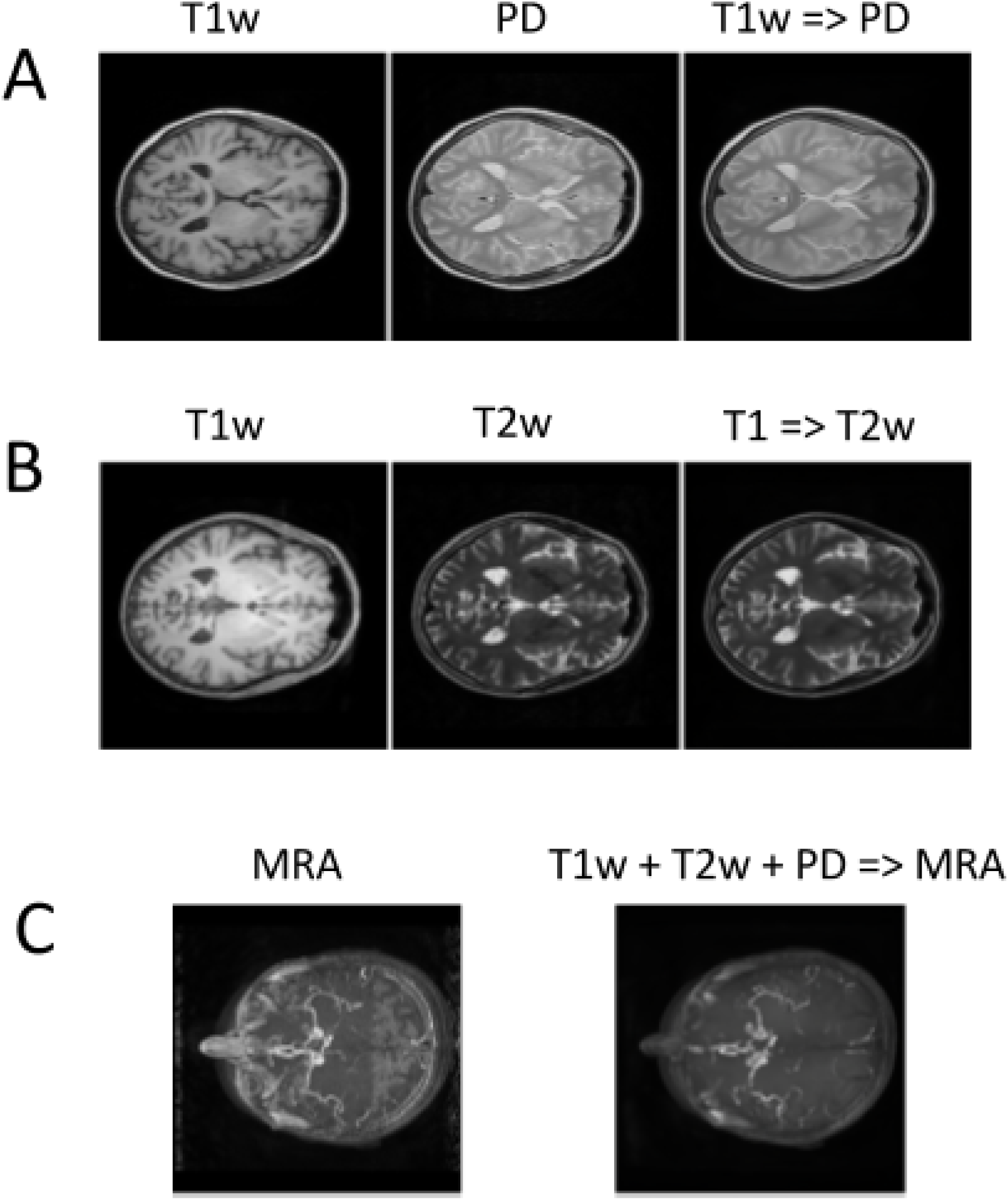
MRI2MRI learns the mapping between different imaging contrasts. (A) Synthesis of PD from T1w: a horizontal slice in one individual brain not shown to the algorithm during training, demonstrating that the algorithm generalizes outside of the training set. Left: input T1w; center: ground truth PD; right: synthesized PD. (B) Synthesis of T2w from T1w: input T1w (left), ground truth T2w (center) and synthesized T2w (right). (C) A many-to-one mapping: combination of T1w, T2w and PD used to synthesize MRA of an individual brain not used during training. The maximum intensity projection along the axial dimension for ground truth (left) and synthesized MRA (right).

The algorithm has practical application in analysis of multi-modal MRI. The algorithm was trained to synthesize T1w images from the DWI b=0 images. On a separate set of subjects, not used during training, we demonstrate that registration of T1w to DWI b=0 is more accurate using a synthesized T1w intermediary. This effect is shown here as mean absolute error (MAE; Figure 2) and is consistent both for the translation component (Mann Whitney U test, p < 10e-7) and for rotation (Mann Whitney U test p<10e-4)

**Figure 2:**
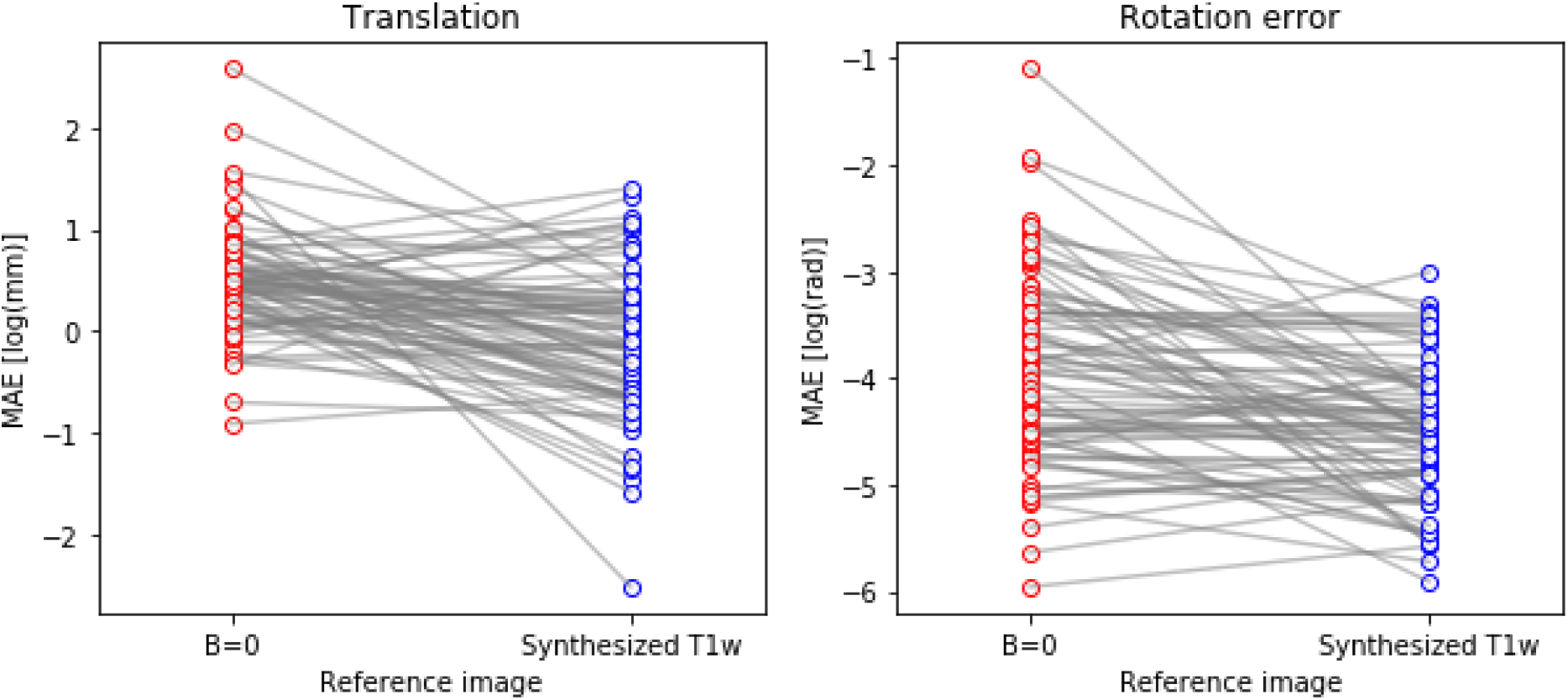
Using images synthesized with MRI2MRI reduces registration errors for registration of DWI data to T1w. In this experiment known affine motion parameters were introduced, and the ability of a state-of-the-art registration algorithm to recover these parameters was assessed, using mutual information as an objective function in registration. We recovered translational (left) and rotational (right) error from the computed registration transformation. These errors are consistently smaller when using the synthesized T1w analog image from DWI b=0 images (y axis in both plots) than when using the b=0 image directly for registration.

## DISCUSSION AND CONCLUSION

We present here a deep learning algorithm that can accurately transform between different MRI contrasts. The algorithm can use as input either one image or more, allowing integration of information from several different contrasts, in generating an image contrast. We present an example where this algorithm can improve the analysis of multi-modal MRI data, by improving the ability to register images with different contrasts to each other, by means of an algorithm-synthesized intermediary image. These results demonstrate that the algorithm is capturing complex nonlinear and spatially-varying dependencies between MRI contrasts, suggesting that it could also be used to better understand these dependencies.

